# MyoView: Fully-automated image segmentation method to analyse skeletal muscle cross section in exercise-induced regenerating myofibers

**DOI:** 10.1101/2021.08.24.457394

**Authors:** Masoud Rahmati, Abdolreza Rashno

## Abstract

Skeletal muscle is an adaptive tissue with the ability to regenerate in response to exercise training. Cross-sectional area (CSA) quantification, as a main parameter to assess muscle regeneration capability, is highly tedious and time-consuming, necessitating an accurate and automated approach to analysis. Although several excellent programs are available to automate analysis of muscle histology, they fail to efficiently and accurately measure CSA in regenerating myofibers in response to exercise training. Here, we have developed a novel fully-automated image segmentation method based on neutrosophic set algorithms to analyse whole skeletal muscle cross sections in exercise-induced regenerating myofibers, referred as MyoView, designed to obtain accurate fiber size and distribution measurements. MyoView provides relatively efficient, accurate, and reliable measurements for detecting different myofibers and CSA quantification in response to the post-exercise regenerating process. We showed that MyoView is comparable with manual quantification. We also showed that MyoView is more accurate and efficient to measure CSA in post-exercise regenerating myofibers as compared with Open-CSAM, MuscleJ, SMASH and MyoVision. Furthermore, we demonstrated that to obtain an accurate CSA quantification of exercise-induced regenerating myofibers, whole muscle cross-section analysis is an essential part, especially for the measurement of different fiber-types. We present MyoView as a new tool for CSA quantification in skeletal muscle from any experimental condition including exercise-induced regenerating myofibers.

## 1. Introduction

Skeletal muscle is an exceptionally regenerative tissue with the ability to undergoes extensive adaptation by changing its fiber type composition and cross-sectional area (CSA) upon external stimuli^1,2^. Exercise training is a unique physiologicall-hypertrophy stimulus with the capability to induce muscle regeneration machinery throw increasing myofiber CSA to overcome corresponding skeletal muscle demands. Moreover, during physical inactivity, aging and some metabolic disorders, skeletal muscle losses in mass due to atrophy of individual myofibers^3^. Among different cellular, molecular and structural components, CSA quantification of myofibers in microscopic images is widely used since it reflects the regenerative capability of the muscle as a final results of activating, proliferating, differentiating and fusing of muscle stem cells^1^. Currently, CSA quantification is commonly performed method to delineate individual myofibers using immunohistochemical approaches targeting laminin or dystrophin in the basal lamina or inside of the sarcolemma, respectively^4,5^. Image quantification is highly time-consuming and labor intensive part of this process and may susceptible to both inter-individual and inter-laboratory variabilities. This is why some laboratories have developed their own automated programs to limit the experimenter bias and save time^4-8^.

Most of current available softwares were developed to measure myofiber CSA in normal muscle or under conditions targeting muscle regeneration including synergist ablation or cardiotoxin injection^4,8^. While, these strategies induce prominent regenerating capability, there are questions about their physiological relevance due to invasive nature and the potential to damage the skeletal muscle^9^. In addition, the shape of the cells in normal muscle is characterized by polygonal and angular myofibers with keeping their contact with each other, while in regenerating myofibers they are round-shaped, highly variable in size, and smallest ones do not regularly contact surrounding fibers. Moreover, image acquisition and reconstitution of the whole muscle section in order to CSA quantification in different fiber types is almost impossible in current available softwares. Depending on the researchers’ decision, different multiple subsets of the whole images will be analysed to find CSAs in different fiber types and it may expose the overall results to bias.

We therefore sought to develop a fully-automated software to quantify CSA in exercise-induced regenerating myofibers. We utilized a high intensity interval training (HIIT) protocol which led to progressive hypertrophy thereby inducing muscle regeneration machinery. Here, we present a fully-automated CSA quantification method for skeletal muscle images applicable to any type of muscle and under exercise-induced regenerating muscle condition. The proposed method; named as MyoView; is based on neutrosophic set algorithms designed to automatically quantify CSA on immunofluorescent picture of the whole skeletal muscle section. In addition, it allows the analysis of the CSAs of different myofibers on the whole muscle cross-section, which we show here to be essential to obtain an accurate CSA quantification.

## 2. Methods

### 2.1. Mice and muscle tissue preparation

All experiments involving animals were performed in accordance with approved guidelines and ethical approval from Lorestan University’s Institutional Animal Care and Use Committee (as registered under the code: LU.ECRA.2017.12). Further, the present study was carried out in compliance with the ARRIVE guidelines. C57BL/6J (n = 18) and mdx (n = 3) mice were purchased from Lorestan University of Medical Sciences Laboratories. At the end of the treatment periods, all mice were anesthetized with inhalation of isoflurane. *Gastrocnemius* muscles from 16- to 18-week-old C57BL/6 mature mice and mdx mice were dissected in optimal cutting temperature (OCT) medium, mounted on pieces of cork, secured with tragacanth gum, frozen in liquid nitrogen-cooled isopentane and stored at − 80 °C. Moreover, samples from regenerating muscles were provided at several time points after exercise training program (day 28 and day 56). Muscle samples were frozen in isopentane cooled by liquid nitrogen and further stored at −80 °C. 10 µm-thick cryosections were prepared and processed for immunostaining and used to test the program’s ability to recognize variability in myofiber morphology.

### 2.2 High-intensity interval training (HIIT) protocol

First, mice were acclimated on the treadmill (5day/week, 10 m/min for 10 minute with no incline) and then subjected to HIIT program for 8 weeks (3 sessions/week). Each training session consisted of a warm-up stage (5 min at 10 m/min), eight exercise intervals at the prescribed speed and angle of inclination for 3–5 min, and a 1 min rest interval at 10 m/min was considered between each interval. The angle of inclination was gradually increased from 10° in the first week to 15° in the second week, 20° in the third week, 25° in the fourth week, and it was maintained at 25° from weeks 4 to 8. The treadmill speed was maintained consistent (15 m/min) for the first 4 weeks and from weeks 5–8 was gradually increased by 1-2 m/min weekly (Model T510E, Diagnostic and Research, Taoyuan, Taiwan)^9^.

### 2.3. Immunofluorescent staining

Immunohistochemical procedures were carried out according to our previous study^10^. In summary, for fiber typing, sections (10 µm-thick) were incubated with antibodies specific to myosin heavy chain (MyHC) types I, IIa, and IIb (BA-D5, SC-71, and BF-F3, respectively, University of Iowa Developmental Studies Hybridoma Bank, Iowa City, IA), supplemented with rabbit polyclonal anti-laminin antibody (L9393; Sigma-Aldrich, St. Louis, MO). MyHC IIx expression was judged from unstained myofibers. Secondary antibodies coupled to Alexa Fluor 405, 488 and 546 were used to detect MyHC types I, IIa, and IIb, respectively (Molecular Probes, Thermo Fisher Scientific, Waltham, MA, USA). Moreover, anti-rabbit IgG Cy3-labeled secondary antibody (Jackson Immunoresearch Labs, West Grove, PA, USA) was used to detect laminin.

### 2.4. Image Acquisition and Quantification

All images were captured at ×10 magnification using a Carl Zeiss AxioImager fluorescent microscope (Carl Zeiss, Jena, Germany). Consecutive fields from whole muscle sections were automatically acquired in multiple channels using the mosaic function in Image M1 Software.

### 2.5. Development of MyoView

MyoView has been implemented in MATLAB 2017b on a machine with 2.26 GHz Corei7 CPU and 8 GB of RAM. It is very fast, simple and efficient with low time complexity to analyze skeletal muscle cross sections. The primary version of source codes is undergoing verification to be publicly available with MIT license at Code Ocean platform in https://codeocean.com/capsule/4910024/tree which generates a standard, secure, and executable research package called a Capsule. Capsule format is open, exportable, reproducible, and interoperable. This capsule is versioned and contains code, data, environment, and associated results of MyoView.

### 2.6. Manual analysis for comparison to MyoView

For manual quantification of fiber-type and CSA, images from various experimental conditions were analyzed in FIJI using the free hand tool to encircle individual myofibers. Manual quantification of CSA and fiber-type were performed for all images used in this present study. Accuracy of MyoView and programs examined in this study was based on comparison between program-derived results and manually acquired results as described^8^. Additionally, Open-CSAM, MuscleJ, SMASH, and MyoVision analyses were performed as described^4,6-8^.

### 2.7. Statistical analyses

Reported data represent mean ± S.E.M. Statistical analysis was performed using the Graph-Pad Prism statistics software (Graph-Pad Software Inc., San Diego, La Jolla CA, USA free demo version 5.04, www.graphpad.com). One-way ANOVA followed by Tukey’s post hoc test was performed for inter-user reliability comparisons. Paired, two-tailed Student’s t-tests were performed for comparing MyoView with manual quantification data. Repeated-measures two-way ANOVA followed by Bonferroni multiple-comparisons tests were performed for CSA changes with HIIT program and fiber counting accuracy and efficiency measurements. Spearman correlation coefficient was computed to assess the correlation analyses.

## 3. Results

### 3.1. Proposed cell segmentation models

#### 3.1.1. Model cell images in neutrosophic sets and neutrosophic images

Interactions between neutralities as well as their scope and nature are modeled in neutrosophy as a branch of philosophy. Neutrosophic logic and neutrosophic set (NS) stem from neutrosophy. Suppose that *N* is a universal set in the neutrosophic domain and a set *X* is included in *N*. Each member *x* in *X* is described with three real standard or nonstandard subsets of [0, 1] named as True(*T), Indeterminacy(I)*, and False(*F)* which have these properties: Sup_T=t_sup, inf_T=t_inf, Sup_I=i_sup, inf_I=i_inf, Sup_F=f_sup, inf_F=f_inf, n-sup=t_sup+i_sup+f_sup and n-inf=t_inf+i_inf+f_inf. Therefore, element *x* in set *X* is expressed as *x(t,i,f)*, where *t, i* and *f* varies in *T, I* and *F* respectively. *x(t,i,f)* could be interpreted as it is *t%* true, *i%* indeterminacy, and *f%* false that *x* belongs to *A. T, I* and *F* could be considered as membership sets [7].

NS can be used in image processing domain. The main contribution of the proposed NS segmentation method is to separate, count and compute sum area of blue, green, black, and red cells in skeletal muscle cross sections. For this task, an image is transformed into the neutrosophic domain. The method of transformation is completely depending on the image processing application. In cell segmentation, image *C* with the dimension of *m×n* and *L* gray levels and *k* channels are considered. Here images with 3 channels Red, Green and Blue (RGB), each channel with the dimension of 5751×7600 for each channel and 256 gray levels are used for automated segmentation. Since all neutrosophic sets are in the range of [0 1], in the first step, C is normalized to interval [0 1] as follows:

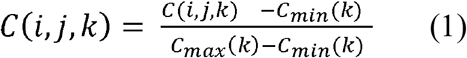

where *C*_*min*_(*k*) and *C*_*max*_(*k*) represent minimum and maximum values of pixels in cell image C in channel k, respectively. *C* is mapped into three sets *T* (true subset), *I* (indeterminate subset) and *F* (false subset). Therefore, the pixel p(i,j) in *C* is transformed into PNS(i, j) = {T(i, j), I(i, j), F(i, j)}) or PNS (t, i, f) in neutrosophic domain. *T, I* and *F* are dedicatedly defined for each type of cells.

#### 3.1.2. Cell counting and area computation for all types of cells

For gray-scale image in Fig. 1A, region of interest is selected and shown in Fig. 1B. The proposed method for cell counting and area computation is explained for this image. In the first step, boundary regions between cells are modeled in true set T and cell regions are considered in False set F. Proposed definitions of neutrosophic sets are as follows:

**Figure 1.**
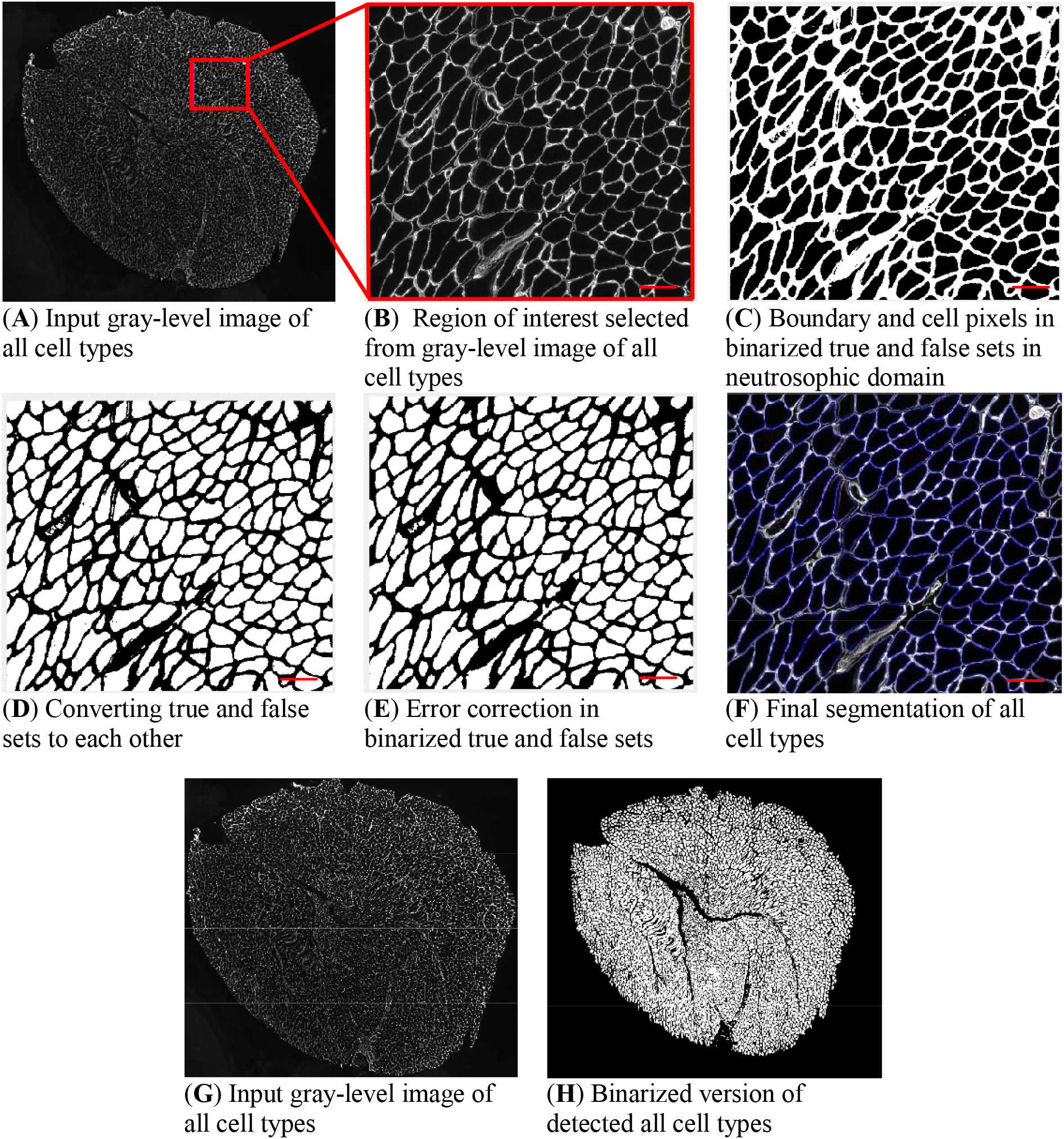
MyoView workflow for all cell types. (**A**) When the MyoView software starts, a window automatically opens to select the image to be analyzed (here muscle cryosections immunostained for laminin in white). (**B**) Represents zoom-in example of a specific area. (**C-F**) Represent MyoView different steps for segmentation process of all cell types. (**G-H**) Represent the segmentation process of all cell types in the whole image. Bars = 25 μm.

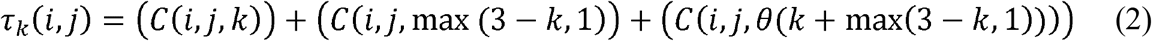

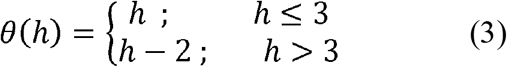

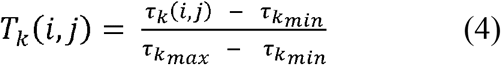

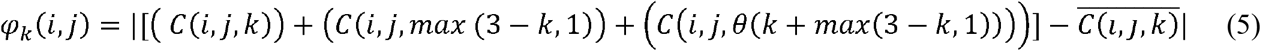

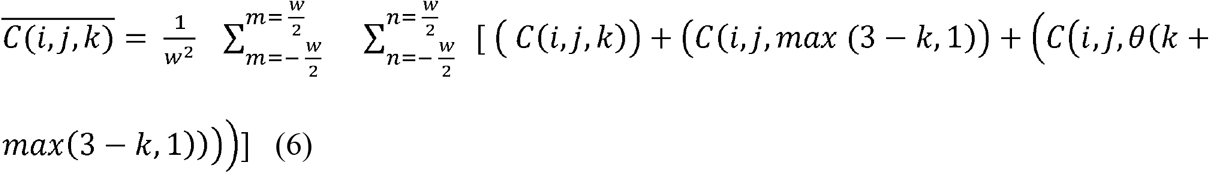

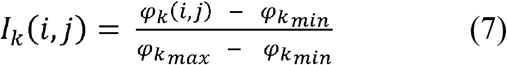

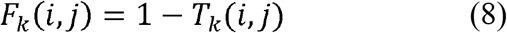

where k is channel number which can be 1, 2 or 3. *min* and *max* indexes are minimum and maximum values in the whole matrix, respectively. Matrix *τ*_*k*_ computes eligibility of pixels to be assigned to cell k, is based on normalized value of pixels in channel k associated with this cell and inverse values in other channels with respect to maximum value 1. True component T is achieved by *τ* normalization. If a pixel has a high value in channel k and low values in other channels simultaneously, a high percent is assigned to this pixel to be a member of cell index k. Indeterminacy matrix is calculated by difference of pixels in channel k from mean of local neighbor pixels in this channel. Therefore, pixels close to local mean of a channel receive low indeterminacy, means a high confidence of assignment is considered for those pixels.

Therefore, in binarized version of True and False sets, boundary and cell pixels are illustrated in light and dark regions, respectively (Fig. 1C). In the next step, true and false sets are converted to each other to place cell pixels in true set as shown in Fig. 1D. In error correction steps, small regions and holes are corrected in Fig. 1E. Finally, true set T is placed in input image and boundaries are illustrated with blue color for better visualization as shown in Fig. 1F. Binarized version of detected all cell types in the whole image is depicted in Figs. 1G-H. In this step, all connected components are found by iteratively 8-neighbor correlated pixels. Components are counted, area of each component is calculated, then number of components, sum and average of areas are reported as outputs.

#### 3.1.3. Cell counting and area computation for color cells

For red, green and blue cells with k indexes of 1,2 and 3, respectively, PNS (t, i, f) means that this pixel is %t percent true to be a member of cell with index k, confidence of this decision is %i and %f percent true that this pixel does not belong to cell k. *T, I* and *F* for cell index k are computed as follows:

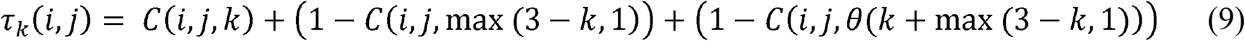

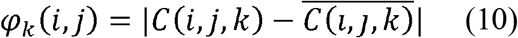

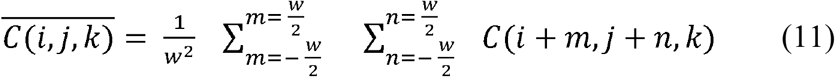

For black cells, Eq. (9) for *τ*_*k*_ computation is rewritten as:

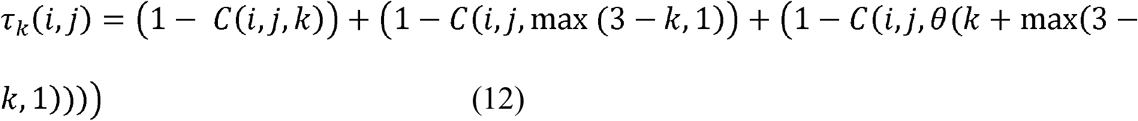

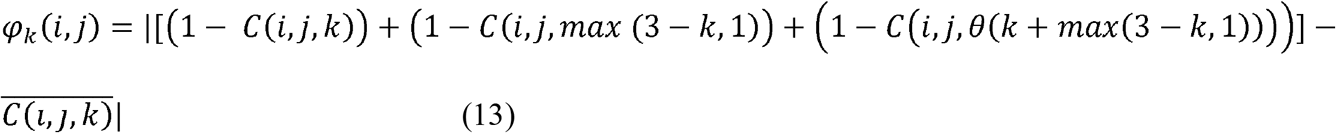

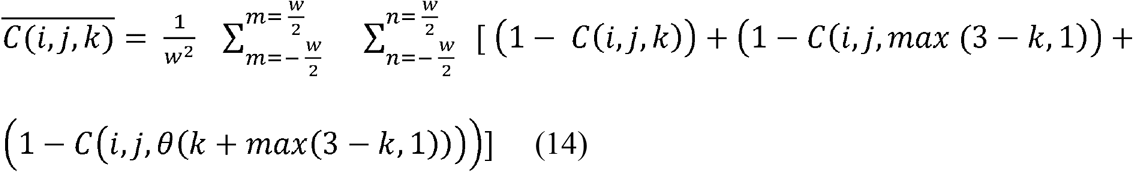

It can be interpreted by this fact that: the lower values of a pixel in all channels, the higher membership degree to black cells is assigned. Consider input image shown in Fig. 2A to apply the proposed segmentation method. For better visualization of details, a region of interest (ROI) is selected and illustrated in Fig. 2B.

**Figure 2.**
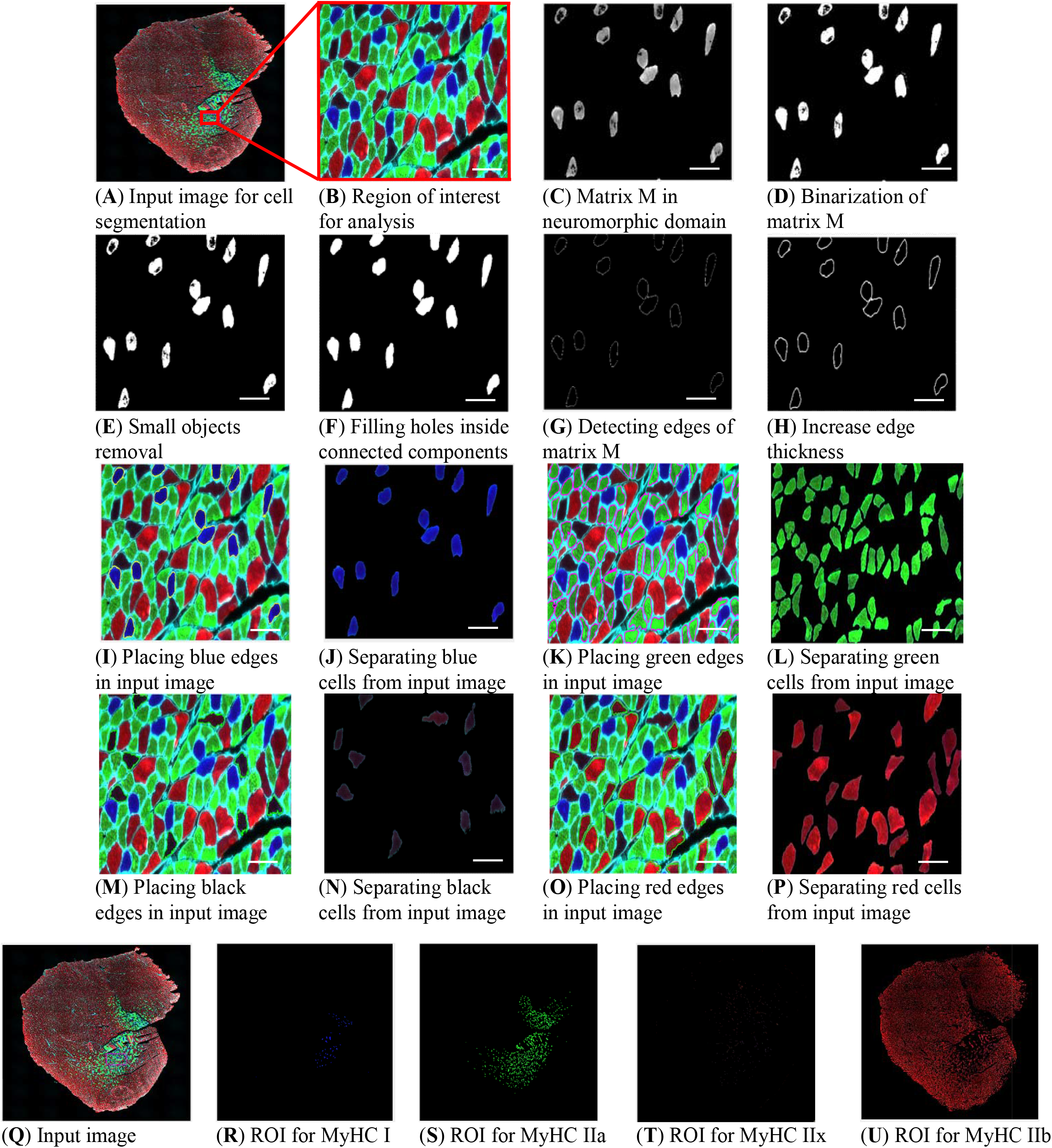
MyoView workflow for different cell types. (**A**) When the MyoView software starts, a window automatically opens to select the image to be analyzed (here muscle cryosections immunostained for laminin (white), MyHC I (blue), MyHC IIa (green), and MyHC IIb (red)). (**B**) Represents zoom-in example of a specific area. (**C**) MyoView computes matrix M in neuromorphic domain from input image. (**D-H**) After binarization of matrix M, removing small objects, and filling holes inside the connected components, then the software detects edges of matrix M and for better visualization increases edge thickness. (**I**) To dabble-check the detecting process, the software places edges in input image. (**J**) Finally, the software separate edges from input image and finalize the analyzing process (here for MyHC I myofibers in blue color). (**K-l**) Represent the same process for MyHC IIa myofibers in green color. (**M-N**) Represent the same process for MyHC IIx myofibers in black color. (**O-P**) Represent the same process for MyHC IIb myofibers in red color. (**Q-U**) Represent the segmentation process of different myofibers in the whole image. Bars = 25 μm.

For each pixel in neutrosophic domain, two conditions are considered to assign high membership degree for that pixel to cell index k. The first one is high value of True matrix and the second one is low indeterminacy, means there is a high confidence to decide that pixel has a high membership degree to True set T. These conditions are combined with “AND” relation by pixelwise product of True and Indeterminacy sets as follows:

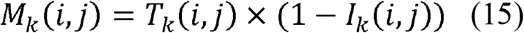

The result of matrix M; still in neuromorphic domain; for blue cells (k=2) is shown in Fig. 2C. It is clear that pixels in blue cells have higher membership degrees (in lighter gray levels) in comparison with pixels in other cells (darker pixels). Matrix M in neuromorphic domain is binarized with a strict threshold as shown in Fig. 2D.

##### 3.1.3.1. Error correction

In binarization process of image M in neutrosophic domain, some extra regions are appeared which are incorrectly assigned to blue regions (Fig. 2D). Therefore, these errors should be corrected. Error correction is done automatically. Connected components for true image T in neutrosophic domain are found iteratively by connecting all 8-neighbor pixels in Supplementary Fig. S1 and labeling connected pixels upon there is no unlabeled pixel. Average area of all components is computed and small components under 20% of average area are ignored as shown in Fig. 2E.

Pixels inside blue cells are located inside a distribution of blue color in channel 3 with a mean and standard deviation. Blue pixels close to the mean of this distribution are strongly assigned to blue cells since high values of T and I matrixes leads to a high value of M for such pixels. Pixels which are far from the mean of distribution are weakly assigned to blue class since their indeterminacy I is high and their true membership is low. Therefore, their values in matrix M are low. It is worth mentioning that such pixels although have low membership degrees blue cells, they are located inside blue regions and should be assigned to blue cells. These errors are corrected by filling holes inside connected components as illustrated in Fig. 2F.

##### 3.1.3.2. Final segmentation of cells

After finalizing matrix M in neutrosophic domain, edge pixels are detected by canny edge detector as shown in Fig. 2G. For better visualization of cell boarders, thickness of edges is increased by image dilation operator with a disk structure element depicted in Fig. 2H. Final edges are placed in input image which lead to high-accuracy segmentation of blue cells in Fig. 2I. Finally, blue cells are separated from input image and shown in an image with black background as illustrated in Fig. 2J. The same scenario is applied to segment red, green and black cells results in detected cells in Figs. 2K-P. Segmented cells in the whole images are shown in Figs. 2Q-U.

### 3.2. MyoView is a reliable software for measuring CSA in response to the post-exercise regenerating situation

In order to test the reliability of MyoView, its performance was compared with some other common software including: Open-CSAM, MuscleJ, SMASH, and MyoVision (Fig. 3). We analyzed *gastrocnemius* muscle from various conditions, including normal muscle, regenerating muscles at several time points after exercise training (D28 and D56) in mature mice, and a model of fibrotic dystrophy (mdx) using anti-laminin antibody. Open-CSAM produced significantly lower mean CSA values as compared with manual quantification on D0, D28 and D56 (Fig. 3A, -6.6, -7.3 and -10.9%, respectively). Mean CSA values obtained with MuscleJ were similar to the manual quantification for normal muscles in D0. However, it gave higher mean CSA values in D28 and D56 post-exercise regenerating muscles (Fig. 3A, between 4.5% to 5.9% of increment). Mean CSA values obtained with SMASH were very close to the manual quantification in normal muscles in D0. However, SMASH produced higher mean CSA values in D28 and D56 post-exercise regenerating muscles (+5.8% and +9.9%, respectively). MyoVision produced similar CSA values to the manual quantification for normal muscles in D0. However, it gave higher mean CSA values in D28 and D56 post-exercise regenerating muscles as compared with manual quantification (Fig. 3A, +9.1% and +8%, respectively). On the other hand, MyoView gave mean CSA values close to those obtained manually in D0, D28 and D56, with a very slight underestimation (Fig. 3A, -1.8%, -1.4% and -1.3%, respectively). In the case of mdx muscle, all of the softwares produced similar CSA values to the manual quantification except MuscleJ and MyoVision which they produced higher values (+4.8% and +9.1%, respectively).

**Figure 3.**
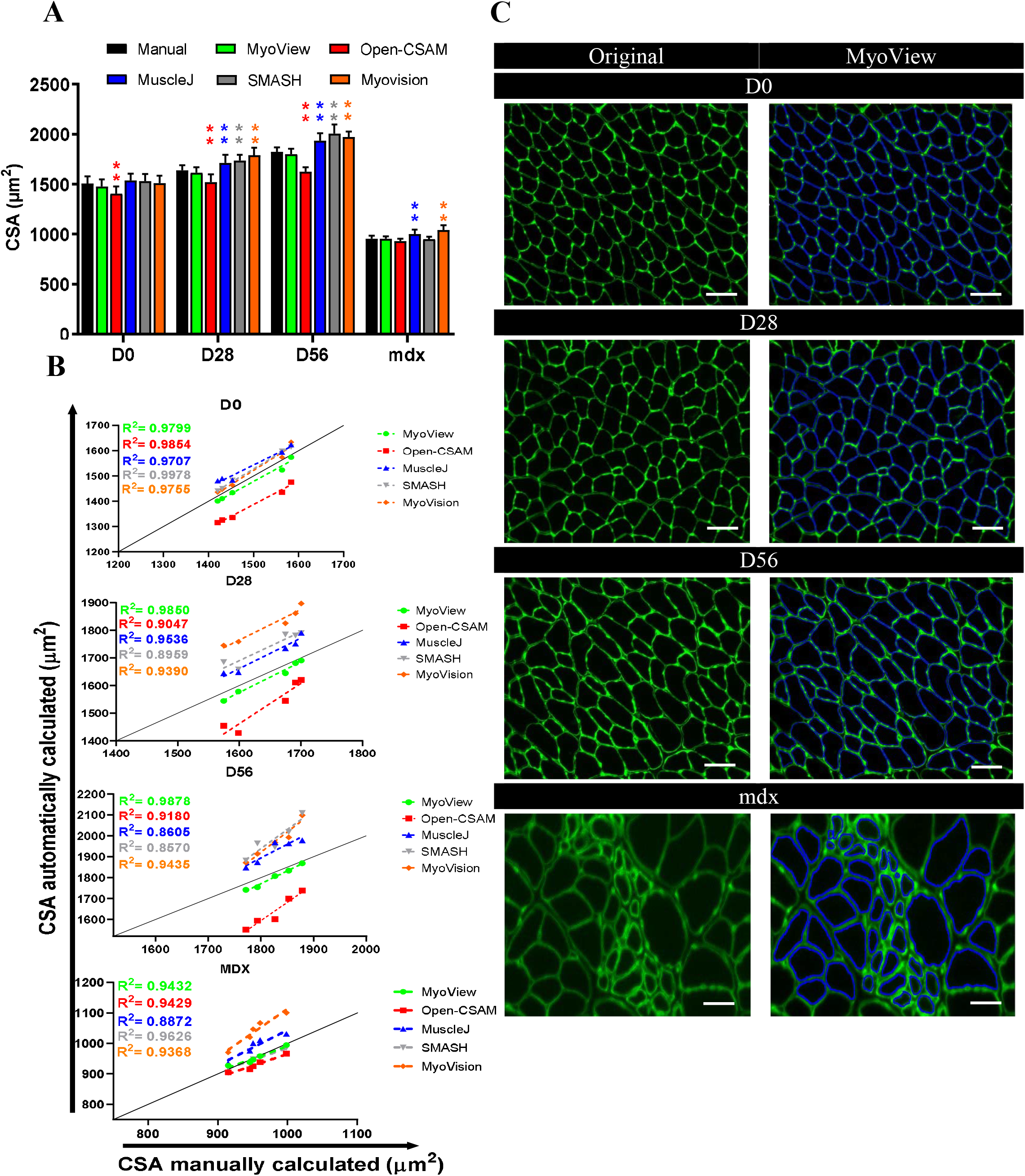
MyoView comparison with Open-CSAM, MuscleJ, SMASH, and MyoVision softwares. The same pictures were analyzed either by manual measurement or using MyoView, Open-CSAM, MuscleJ, SMASH, and MyoVision softwares. (**A**) Mean cross-section area (CSA) obtained on various *gastrocnemius* muscles. Muscles were isolated from 16- to 18-week-old C57BL/6 mice in (D0) or 28 days (D28), and 56 days (D56) post-HIIT program, and from dystrophic mice (mdx). Results are mean ± SEM of 10 images from 5 muscles in each conditions. (**B**) Correlation between manual measurement (X axis) and MyoView, Open-CSAM, MuscleJ, SMASH, and MyoVision softwares (Y axis) measurements performed on the same images used in (**A**). (**C**) Representative images measured by MyoView on days 0, 28, and 56 post-HIIT program, and from dystrophic mice (mdx). Bars = 25 μm. **p < 0.01 as compared with manual quantification.

Despite an increased CSA values obtained using MuscleJ, SMASH, and MyoVision and decreased values for Open-CSAM, the correlation between them and manual quantifications were strong in normal muscles in D0, as well as in days 28 and 56 post-exercise regenerating muscles (Fig. 3B, R^2^ > 0.85), suggesting that in these conditions, CSA overestimation by MuscleJ, SMASH, and MyoVision as well as CSA underestimation by Open-CSAM were similar to all the pictures and did not introduce a specific bias. Overall correlation between Open-CSAM and manual quantification was very strong in normal muscles in D0 (Fig. 3B, R^2^ = 0.9854). Although this correlation was lower in days 28 and 56 post-exercise regenerating muscles as well as on fibrotic muscles (R^2^ = 0.9047, R^2^ = 0.9180, and R^2^ = 0.9429, respectively). Similarly, overall correlation between MuscleJ, SMASH, and MyoVision and manual quantification were very strong in normal muscles in D0 (Fig. 3B, R^2^ = 0.9707, R^2^ = 0.9978, and R^2^ = 0.9755, respectively). Although these correlations were lower in D28 (Fig. 3B, R^2^ = 0.9536, R^2^ = 0.8959, and R^2^ = 0.9390, respectively) and D56 post-exercise regenerating muscles (Fig. 3B, R^2^ = 0.8605, R^2^ = 0.8570, and R^2^ = 0.9435, respectively) as well as on fibrotic muscles (R^2^ = 0.8872, R^2^ = 0.9626, and R^2^ = 0.9368, respectively). Although, the correlation between MyoView and manual quantification was very strong in normal muscles in D0 (Fig. 3B, R^2^ = 0.9799), there was no difference between this correlation and the corresponding values for other software. However, the correlation between MyoView and manual quantification was better than Open-CSAM, MuscleJ, SMASH, and MyoVision in 28 and 56 days’ post-exercise regenerating muscles. This suggests that MyoView performance was better in response to the post-exercise regenerating process.

### 3.3. MyoView is an efficient and accurate software for measuring CSA

In order to examine MyoView efficiency in detecting myofibers and the time spent on CSA analysis as well as its accuracy, we next compared MyoView performance with manual quantification, Open-CSAM, MuscleJ, SMASH, and MyoVision softwares (Fig. 4). Figure 4B shows that there was no difference between the number of fibers counted by MyoView and the number counted by manual quantification, which corresponds to an accuracy of 98.1% ± 0.9 (Fig. 4D). In contrast, Open-CSAM, MuscleJ, SMASH, and MyoVision identified lower myofibers and spent much more time to analyse CSA from various experimental conditions (Fig. 4C, P > 0.001). Moreover, as compared with manual quantification, the accuracy of MyoView in analysing CSA was 98.2% ± 1.4, (Fig. 4E), while Open-CSAM, MuscleJ, SMASH, and MyoVision have been less accurate in analysing CSA (P > 0.001). Taken together, these results suggest that MyoView is an efficient and accurate software for detecting myofibers and measuring CSA in response to the post-exercise regeneration process.

**Figure 4.**
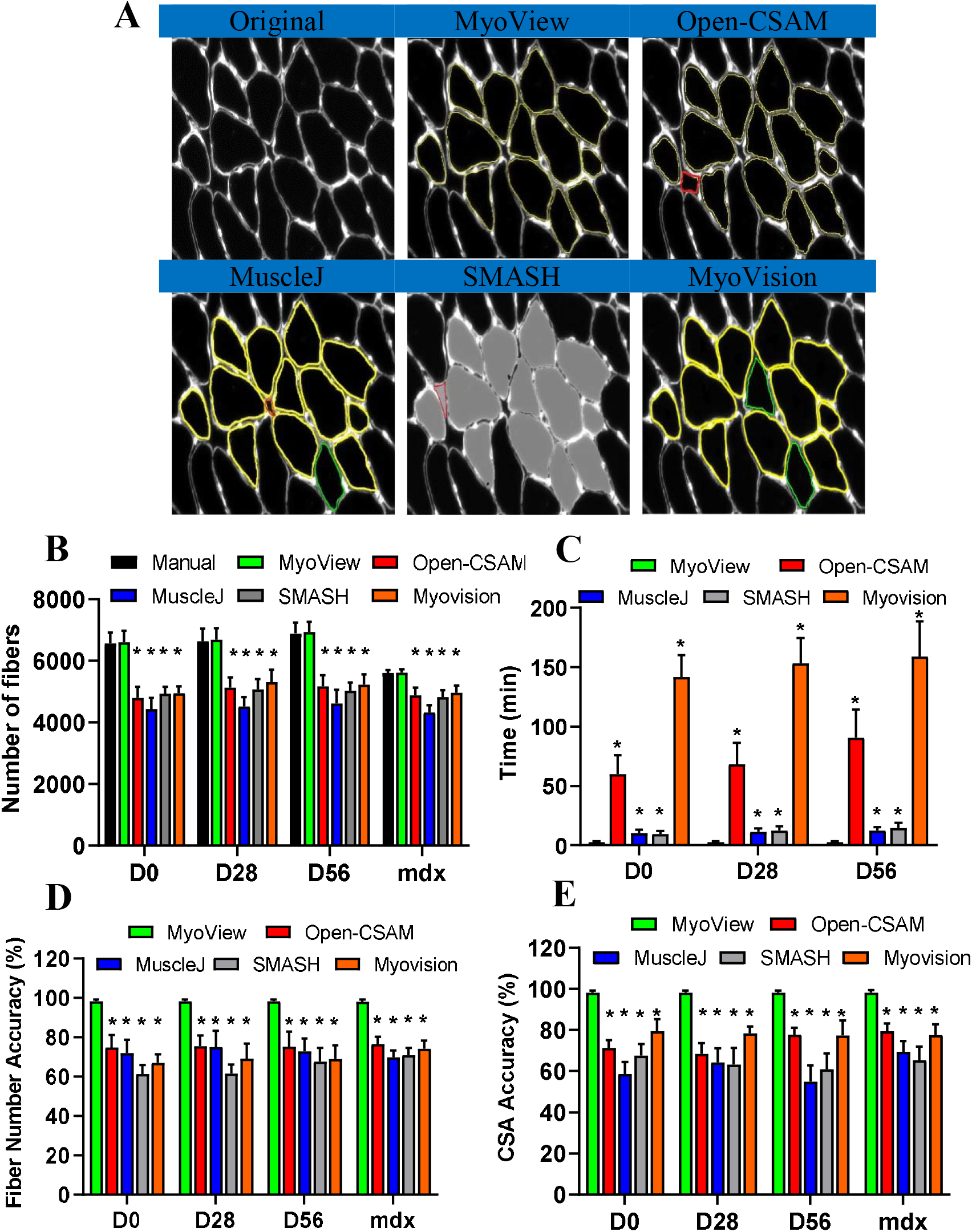
Comparison of MyoView, Open-CSAM, MuscleJ, SMASH, and MyoVision CSA quantification on whole muscle sections. (**A**) Representative images measured by MyoView, Open-CSAM, MuscleJ, SMASH, and MyoVision on *gastrocnemius* muscle images obtained from 16- to 18-week-old C57BL/6 mice. Segmentation errors are labeled as missed fibers (green), mis-segmented fibers (red). (**B**) Number of fibers identified by the softwares. (**C**) Total analysis time required by the different softwares. (**D**) Fiber number accuracy by the different softwares. (**E**) Mean CSA accuracy by the different softwares. Results are mean ± SEM of 10 images from 5 muscles in each condition. **p < 0.01 as compared with manual or MyoView quantification.

### 3.4. MyoView performance in different fiber-types is comparable to manual quantification

We next wanted to determine how does effective MyoView work as a tool for analysing different myofiber size and type in entire cross-section of *gastrocnemius* muscle. Three experienced researchers used the free hand tool in Fiji to encircle individual myofibers from six images from 16-to 18-week-old C57BL/6 mice in 56 days (D56) post-HIIT program to obtain CSA values. We then ran the same images through the MyoView program and obtained a distribution of CSA across the images. The mean CSAs and distributions did not differ significantly between manual and MyoView analysis (Fig 5A-B). Next, we tested the accuracy of fiber typing using MyoView. Fiber type analysis was manually performed by 3 experienced researchers on six images (2 images per person). We then used MyoView to obtain mean data for blue, green, black, and red channels across these six images for fiber typing. The relative proportion of each fiber type was strongly correlated between MyoView and manual analysis (R^2^ > 0.97) (Fig 5C-D). Additionally, MyoView fiber type classification results in CSA were linearly and positively correlated to manual counts (R^2^ > 0.98), and there was no statistically significant difference between the CSAs of each fiber type measured by hand and by MyoView. The accuracy of MyoView fiber type analysis is estimated to be 98.5 ± 0.7% compared with manual quantification. Taken together, the results from this part implicate that MyoView performance in different fiber-types is comparable to manual quantification in regenerating myofibers in response to HIIT program.

**Figure 5.**
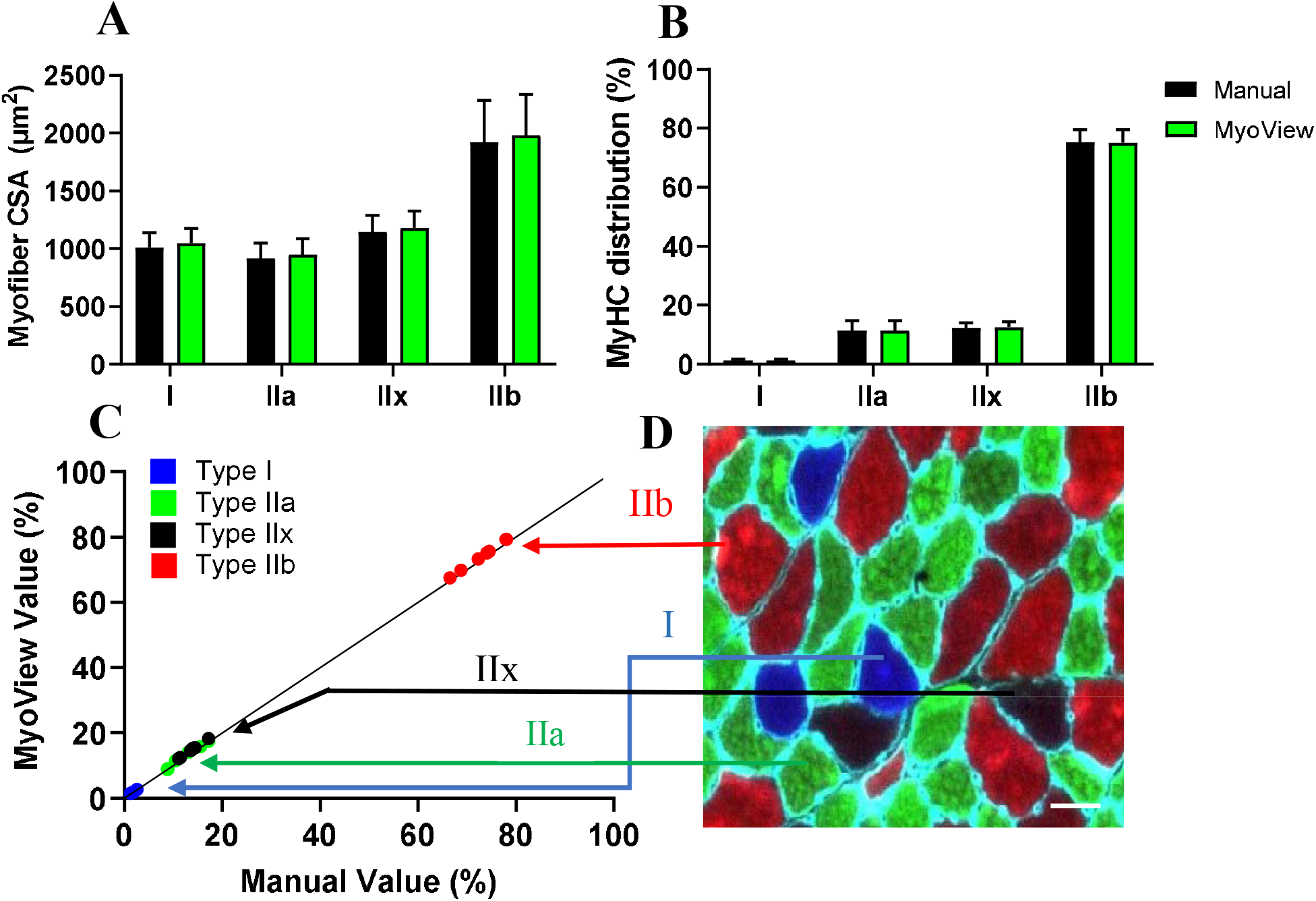
MyoView is comparable to manual quantification for analysing different myofibers. (**A-B**) Myofiber CSAs and distributions were not different when determined with MyoView or by manual quantification (p>0.92). Results of manual analysis of 6 images from 3 investigators are shown. (**C**) Proportions determined manually are on the y-axis and proportions determined by MyoView are on the x-axis. (**D**) MyHC I fibers indicated by blue symbols, MyHC IIa indicated by green symbols, MyHC IIx indicated by black symbols, and MyHC IIb indicated by red. Bar = 25 μm.

### 3.5. Inter-user reliability of MyoView

To assess the ease and accuracy of analyses with MyoView, we asked five individuals in the laboratory to analyze the CSAs of whole muscle cross-section and different fiber-types from a single image from *gastrocnemius* muscle from D28 after exercise training. The analyses of the CSA of whole muscle cross-section, MyHC I, IIa, IIx, and IIb fibers were similar among the five users (Fig. 6). This further demonstrates the reliability of the image outputs of the analyses taken by MyoView to analyses the CSAs of whole muscle cross-section and different fiber-types in regenerating myofibers in response to HIIT program.

**Figure 6.**
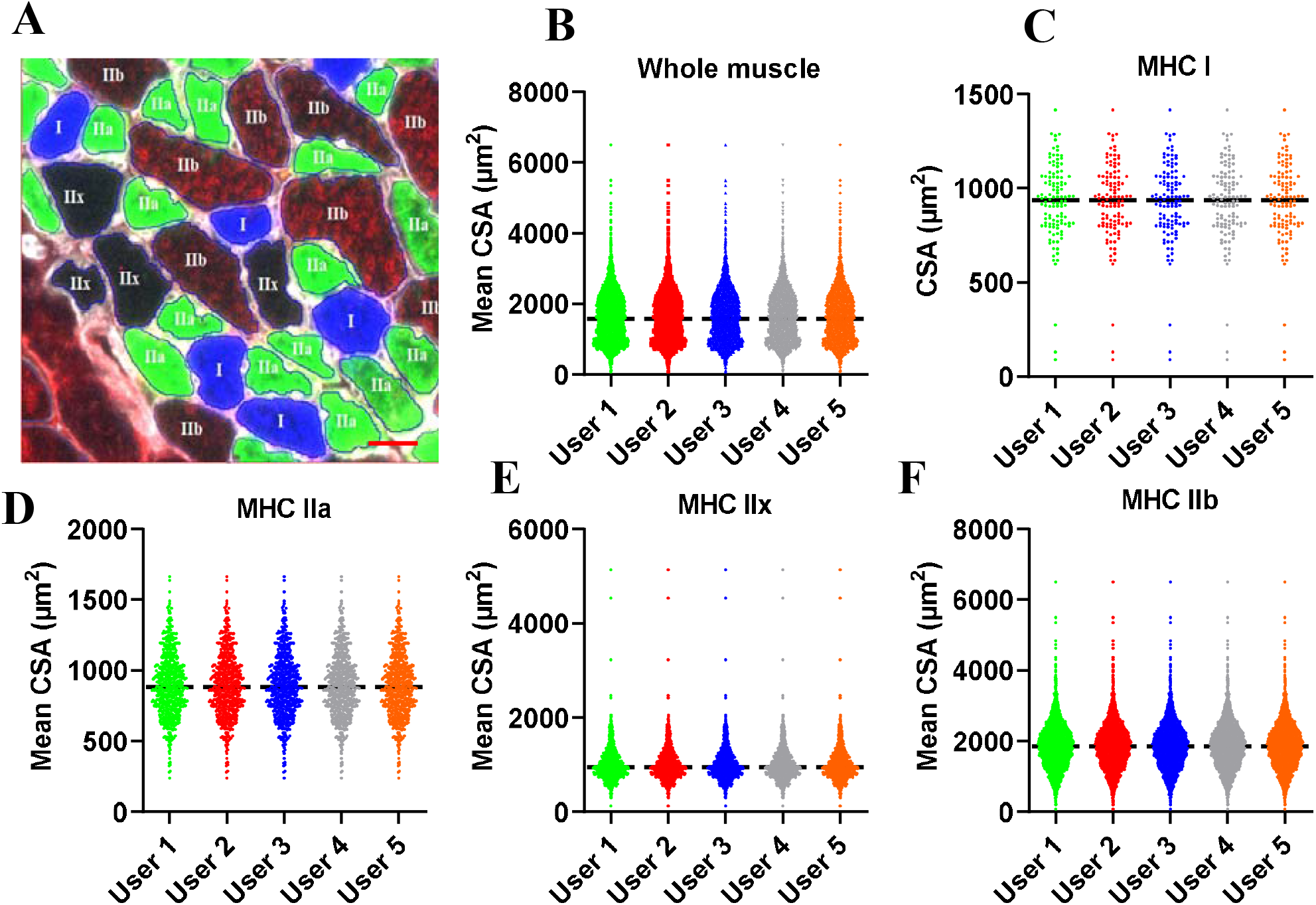
Inter-user reliability. Independent analyses of mean CSA as a function of fiber type by 5 users. Data from whole *gastrocnemius* muscle images obtained from 16- to 18-week-old C57BL/6 mice on 56 days (D56) post-HIIT program. (**A**) Representative image measured by MyoView for measuring different fiber types. (**B**) Mean CSA of whole muscle fibers. (**C**) Mean CSA of MyHC I fibers. (**D**) Mean CSA of MyHC IIa fibers. (**E**) Mean CSA of MyHC IIx fibers. (**F**) Mean CSA of MyHC IIb fibers. Bar = 25 μm.

### 3.6. Whole muscle cross-section analysis for fiber type determination is essential for best accuracy

CSA determination of the of various fiber types is usually performed on a subset of images randomly taken throughout the muscle section. Depending on the researchers’ decision, a variety of different number of images and thus myofibers can be qualified for fiber type analysis. This may expose the evaluation process to the possibility of selection bias as myofiber size is quite heterogeneous through the whole muscle cross-section. Figure 7A shows an example of an entire reconstituted muscle picture. We measured CSAs of different myofibers on individual images from 5 mice on 56 days post-HIIT program, calculated the mean CSAs on 12 subsets of images, and compared the results with the CSAs obtained on the whole muscle section by MyoView. When the measurement was made only using a 12 subset of pictures, there was no significant different in fiber type distribution as compared with whole muscle cross-section analysis (Fig. 7B, P > 0.8). Given that about eighty percent of *gastrocnemius* muscle fibers are type IIb, measuring fewer number of these fibers on 12 subsets of pictures (Fig. 7C, P = 0.015) was led to underestimation of CSA of IIb fibers as compared with whole muscle cross-section analysis (Fig. 7D, P > 0.8). Moreover, compared with whole muscle cross-section analysis, significant reduced CSA of type IIx fibers was observed in analyzing of 12 subsets of pictures (Fig. 7D, P = 0.04). Additionally, we particularly observed that compared with whole muscle cross-section analysis, mean CSA was underestimated when 12 subsets of pictures were measured (Fig. 7C, P = 0.015). Taken together, these results indicate that the whole muscle cross-section should be analyzed when measuring CSA of exercise-induced regenerating muscle in order to obtain an unbiased data.

**Figure 7.**
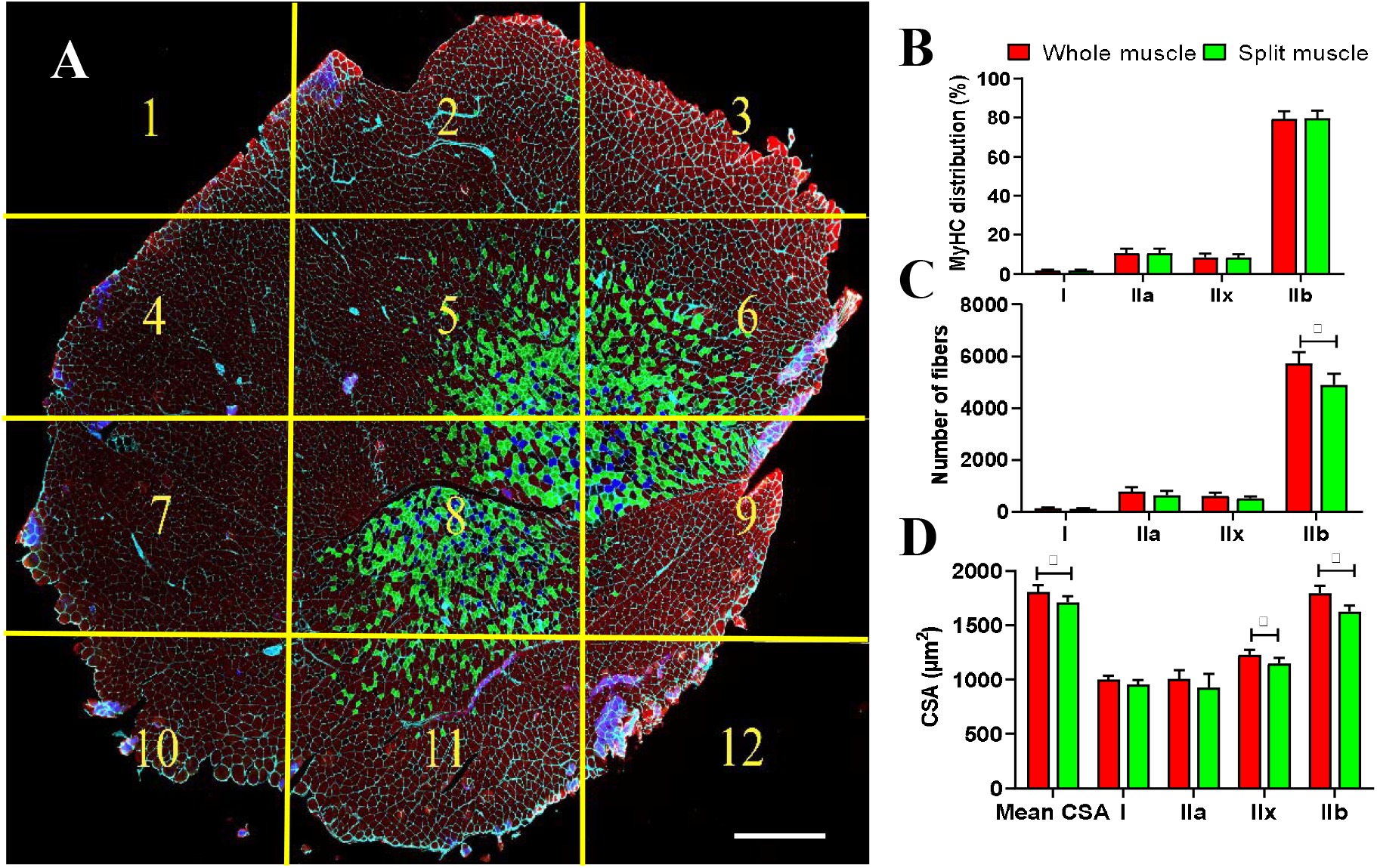
Whole muscle analysis for different myofiber types by MyoView. (**A**) Whole reconstitution of a MyHC I, IIa, IIx, and IIb -stained cryosection of a *gastrocnemius* muscle from 5 mice on 56 days post-HIIT program (12 pictures were automatically recorded and assembled by MetaMorph software). The position of each individual image is highlighted by the yellow lines. (**B**) MyHC distribution obtained after various subsettings of pictures in (**A**) as compared with whole muscle analysis by MyoView. (**C**) Number of fibers obtained after various subsettings of pictures in in (**A**) as compared with whole muscle analysis by MyoView. (**B**) Mean cross-section area obtained after various subsettings of pictures in in (**A**) as compared with whole muscle analysis by MyoView. Bar = 250 μm.

## 4. Discussion

Skeletal muscle fibers are extremely sensitive to exercise training stimuli, with individual myofibers capable to increase in size^9,10^. Due to this adaptive characteristic, exercise physiologists have long acknowledged the importance of accurately quantifying muscle CSA in their experiments^8^. However, there are various strategies to analyze images and CSA quantification that can give highly heterogeneous results among different laboratories and teams. On the other hand, no automated program has been developed to provides the possibility for CSA quantification in exercise-induced regenerating myofibers especially in whole muscle cross-section. In the present study, we presented MyoView software to automatically process immunofluorescence images of the whole muscle cross-sections stained with laminin α2 and antibodies specific to MyHC types I, IIa, and IIb (BA-D5, SC-71, and BF-F3, respectively) in order to facilitate the determination of different individual myofibers on D0, D28 and D56 post-exercise training. The parallel comparison between MyoView and manual quantification showed that MyoView can provide relatively efficient, accurate, and reliable measurements for detecting different myofibers and measuring CSA in response to the post-exercise regeneration process.

MyoView is based on neutrosophic set algorithms. It is a fully-automated method for color cell segmentation based on neutrosophic sets. To the best of our knowledge, this is the first method which is proposed for neutrosophic cell segmentation. The main benefit behind using neutrosophic set is that: it has been applied in many applications; including segmentation of fluid/cyst regions in diabetic macular edema and exudative age-related macular degeneration patients^11-13^, unsupervised color–texture image segmentation^14^, automatic segmentation of choroid layer in retinal images^15^, content-based image retrieval^16,17^, and promising results were achieved. In this research, first, color cells have been modeled as neutrosophic sets with three components and then each component is used to increase the confidence of each pixel to its corresponding cell type. Therefore, a high-confidence assignment of pixels to cell regions is achieved.

Several other semi and fully-automated softwares have been developed^4-8,18-25^. Among them, we have attempt to test and compare Open-CSAM, MuscleJ, SMASH, and MyoVision with MyoView and found MyoView is relatively easy to implement and more accurate for CSA quantification, especially in post-exercise regenerating muscles. We did not test all available softwares as they are either not available online or purchase is required. Open-CSAM, MuscleJ, SMASH, and MyoVision are well-designed software packages that act as free versions of commercially available image analysis tools, but they require varying amounts of manual corrections to ensure accuracy, especially when it comes to analysing different fiber type across the muscle section. The primary goal of MyoView was to develop an accurate fully-automated software for whole muscle cross-section that is user friendly and requires minimal post-analysis corrections. The accuracy in the CSA quantification and identifying different fiber types are enhanced by MyoView, especially in regenerating muscles in response to exercise training stimuli.

There are several limitations for the current version of our MyoView software. First, the accuracy of CSA quantification depends on the quality of the immunostaing procedures. In this case, we recommend performing a new immunostaing rather than trying to analyze poor quality images. Second, MyoView does not allow to manually correct the wrongly identified muscle fibers. However, our initial study with the 100 mages showed that MyoView error for myofiber identification was less than 3% which it does not appear to affect the conclusion of CSA quantification. Finally, in the current version of MyoView we have not provided the possibility to count and analyze myonuclei, vessels, and satellite cells. Future improvements can be made to develop these functions in MyoView.

We present a new fully-automated image analysis program, MyoView, for analyses of whole muscle cross-sectional area and fiber-type distribution in exercise-induced regenerating myofibers. MyoView allows rapid and accurate analysis of whole muscle cross-sectional immunofluorescence images. Additionally, MyoView rapidly identifies different myofibers based on the expression of myosin heavy chain isoforms in skeletal muscle from any experimental condition including exercise-induced regenerating myofibers.

## Author Contributions

M.R. and A.R. conceptualized and designed the research; M.R. performed the data analysis and interpretation; A.R. developed the software; M.R. and A.R. wrote the article.

## Acknowledgements

This research was supported by the Lorestan University through grant number 99101418.

